# Global SLAM-Seq for accurate mRNA decay determination and identification of NMD targets

**DOI:** 10.1101/2021.12.20.473500

**Authors:** Hanna Alalam, Jorge Zepeda, Per Sunnerhagen

## Abstract

Gene expression analysis requires accurate measurements of global RNA degradation rates, earlier problematic with methods disruptive to cell physiology. Recently, metabolic RNA labeling emerged as an efficient and minimally invasive technique applied in mammalian cells. Here, we have adapted SH-Linked Alkylation for the Metabolic Sequencing of RNA (SLAM-Seq) for a global mRNA stability study in yeast using 4-thiouracil pulse-chase labeling. We assign high-confidence half-life estimates for 67.5 % of expressed ORFs, and measure a median half-life of 9.4 min. For mRNAs where half-life estimates exist in the literature, their ranking order was in good agreement with previous data, indicating that SLAM-Seq efficiently classifies stable and unstable transcripts.

We then leveraged our yeast protocol to identify targets of the Nonsense-mediated decay (NMD) pathway. There are currently no global reports of half-lives in both wild type and NMD defective yeast cells; instead steady-state RNA level changes are used as a proxy. With SLAM-Seq, we assign 580 transcripts as putative NMD targets, based on their measured half-lives in wild-type and *upf3Δ* mutants. We find 230 novel targets, and observe a strong agreement with previous reports of NMD targets, 60 % of our candidates being identified in previous studies.

This indicates that SLAM-Seq is a simpler and more economic method for global quantification of mRNA half-lives. Our adaptation for yeast yielded global quantitative measures of the NMD effect on transcript half-lives, high correlation with RNA half-lives measured previously with more technically challenging protocols, and identification of novel NMD regulated transcripts that escaped prior detection.

## INTRODUCTION

Post-transcriptional controls make major quantitative contributions to regulation of gene expression. These include regulation of the decay and translation rates of an mRNA species, as well as its splicing and intracellular location. The level of an RNA species is determined by its rates of synthesis and degradation. While control of transcriptional initiation has been extensively studied, understanding of RNA decay is lagging behind. Turnover rates of RNA molecules are intimately linked to other post-transcriptional processes, such as translation (Pelechano et al. 2015; Chan et al. 2018; Hanson and Coller 2018), intracellular localization (Bovaird et al. 2018), and sequestering in RNA granules (Sheth and Parker 2003; Huch et al. 2016; Escalante and Gasch 2021). Established methods to quantitate the decay rate of RNA species *in vivo* have limitations that hamper progress in this regard. The stability of individual mRNAs can be reliably quantitated in low throughput by placing them under control of a regulatable promoter, and measuring half-life after transcriptional shut-off (Baudrimont et al. 2017). For global analyses, RNA synthesis can be arrested by inhibition of RNA polymerases, and the degradation rate of all RNA species monitored alternatively metabolic labeling approaches can be used. The budding yeast *Saccharomyces cerevisiae* has been at the forefront of developing technologies to study most aspects of post-transcriptional regulation, by virtue of its genetic tractability and ease of performing genome-wide studies. In *S. cerevisiae*, several global studies of mRNA decay rates have been performed. Beyond the basal rates, the effects on global mRNA degradation by environmental stress (Molin et al. 2009; Romero-Santacreu et al. 2009; Miller et al. 2018) or defects in RNA decay pathways (Guan et al. 2006; Celik et al. 2017) have been studied. In *S. cerevisiae*, a heat-sensitive allele of the largest subunit of RNA polymerase II, *rpb1-1*, can be used to block mRNA synthesis at the restrictive temperature (Nonet et al. 1987). The downside of this approach is that temperature upshift activates heat shock responses and changes the physiology of the cell. Chemical RNA polymerase II inhibitors, 1,10-phenanthroline, thiolutin, and 6-azauracil, have likewise been used for this purpose. Again, complications arise since each of them selectively induces transcription of specific gene groups (Grigull et al. 2004; Eshleman et al. 2020), confounding their analysis. Both for chemical and genetic blockade of RNA synthesis, perturbations of the physiological state of the cell are thus problematic. These weaknesses of the methods are enhanced when studying transient phenomena such as changes in mRNA stability under environmental perturbations (Rodríguez-Gabriel et al. 2006; Molin et al. 2009; Romero-Santacreu et al. 2009). First, the impact of the heat shock or the chemical inhibitor themselves will obscure that from the environmental perturbation; second, the resolution in time is poor and comparable to the mRNA half-life itself and to the time scale of its changes on environmental shocks.

Attempting to avoid perturbations from the method of study itself, decay rates for RNA species can be indirectly calculated from data on their synthesis rates and steadystate levels, *e.g*. from genomic run-on experiments using *in vivo* radioactive labeling (Marín-Navarro et al. 2011; Jordán-Pla et al. 2019). More recently, metabolic labeling of RNA using nucleoside analogs such as 4-thiouridine (4sU) has been employed to study the transcriptional process (Russo et al. 2017). Alternatively, other nucleoside analogs including 5–bromouridine (Tani et al. 2012) or 5–ethynyluridine (Abe et al. 2012) can be used, nevertheless studies in mammalian cells have relied mostly on 4sU. This nucleoside analog cannot be imported into unmodified yeast cells, however. On the other hand, the enzyme uracil phosphoribosyltransferase will metabolize the base analog 4-thiouracil (4tU) into the nucleotide analog 4-thiouridine monophosphate, allowing its incorporation into nascent RNA in yeast, but not in mammalian cells. Consequently, 4tU is instead the preferred labeling choice for budding or fission yeast.

The thiolation of 4tU enables it to engage in reversible covalent conjugation to thiol-containing molecules such as streptavidin. This can be employed for biochemical enrichment using solid phase immobilization of thiolated mRNAs. With pulse-chase labeling protocols, this allows detailed kinetic studies of transcription and RNA degradation. Such protocols have been used for global measurements of transcript stability in yeast (Chan et al. 2018). In that work, the mean half-life of *S. cerevisiae* mRNA was estimated to be under 5 min, considerably lower than in previous studies. A mean value of 23 min was measured using temperature shift with *rpb1-1* (Wang et al. 2002), and a mean of 35.3 min was calculated from experiments where tagged Rbp1 was depleted from the cell nucleus (Geisberg et al. 2014). One possible reason for such large discrepancies between half-life estimates using inhibition of RNA synthesis versus using metabolic labeling is that RNA polymerase inhibition may also affect RNA degradation. It was observed that application of thiolutin to yeast cells increased apparent mRNA half-life in a dose-dependent manner (Pelechano and Pérez-Ortín 2008). It has been proposed that the processes of mRNA synthesis and degradation promote each other, and that mRNA synthesis inhibition leads to reduced mRNA decay and *vice versa* (Haimovich et al. 2013). In a further development of metabolic labeling (“SLAM-Seq”), after extraction of RNA, the incorporated thiolated uracil derivatives are instead alkylated with iodoacetamide (Herzog et al. 2017). In the subsequent reverse transcription reaction, guanine will base pair with the iodoacetamide uracil derivative, resulting in a T-to-C conversion in the position originally occupied by 4tU. This eliminates errors and time consumption from the biochemical purification steps.

Nonsense-mediated decay (NMD) was originally discovered in *S. cerevisiae* through the accelerated degradation of mRNAs with premature stop codons (PTCs) (Losson and Lacroute 1979). The sources of PTCs can be transcriptional error, defective splicing, or translational shifting. It has subsequently emerged that NMD can be triggered by other structures than PTCs, and affects a large proportion of mRNAs. Thus, long 3’-untranslated regions (3’-UTRs) can also bring about NMD (Hogg and Goff 2010; Hurt et al. 2013). It has been calculated that up to 20 % of all eukaryotic RNAs are affected directly or indirectly by NMD (Mendell et al. 2004; Guan et al. 2006). The core proteins required for NMD are Upf1 – 3, present throughout eukaryotes and first discovered in *S. cerevisiae* (Leeds et al. 1992). Among these, the RNA helicase Upf1 is the most highly conserved in evolution. The Upf2 and Upf3 proteins are thought to bridge the effector Upf1 and the structure triggering the NMD reaction, stimulating its helicase activity (Chamieh et al. 2008). It has been proposed that the Upf proteins are also required for processes beyond NMD, *e.g*. telomere maintenance, through incompletely elucidated mechanisms (Askree et al. 2004). On the other hand, these effects may also be mediated through telomere maintenance mRNAs being NMD targets. In metazoans, the SMG1 protein kinase activates Upf1 through phosphorylation; yeast Upf1 also becomes phosphorylated but its kinase is as yet unidentified (Lasalde et al. 2014).

The total population of NMD targets is not fully characterized in any organism. Such a characterization is an extensive task, as many RNA species arise from one transcriptional unit, and these can have different propensity to be targeted by NMD. Moreover, the extent to which a particular RNA species is degraded by NMD may vary between conditions, as NMD is also regulated by *e.g*. stress conditions (Goetz and Wilkinson 2017). By measuring steady-state levels of *S. cerevisiae* mRNA species in wt and *upf* mutants, direct and indirect NMD targets were found to comprise around 15 % of the transcriptome (Celik et al. 2017).

Here, we applied SLAM-Seq for *S. cerevisiae* using 4tU; previously, this protocol was optimized for mammalian cells using 4sU. We have determined individual mRNA half-lives in *S. cerevisiae* cells in the BY4741 background and compared wild-type (wt) with the *upf3Δ* mutant, defective in NMD. We provide the first application of SLAM-Seq to study mRNA stability in yeast. Further, we assign a direct quantitative measure of the change of mRNA half-life in an NMD defective background, rather than only observing changes in RNA abundance, which also includes indirect effects through *e.g*. mRNAs encoding transcription factors as NMD targets. SLAM-Seq obviates the need for biochemical purification of derivatized mRNA by utilizing high efficiency chemical conversion. We paired SLAM-Seq with 3’-UTR end sequencing after having scrutinized the annotation of 3’-UTRs genome-wide to allow cost effective assessment of the stability of polyadenylated mRNA transcripts (Herzog et al. 2020), and capture a total of 3033 transcripts for which we can assign a half-life estimate with high precision in both the wt and the NMD mutant in biological triplicate. We find an average mRNA half-life of 11 min (median 9.4 min) in wt and 13 min (median 11.6 min) in the *upf3Δ* mutant. We identify transcripts from 580 genes as being under the control of NMD under standard conditions. These include 350 transcripts that have been found in earlier studies, and 230 not previously recognized.

## RESULTS

SLAM-Seq utilizes a highly efficient derivatization of thiol-containing uridines with iodoacetamide that eventually leads to specific mis-incorporation of a cytosine (C) instead of a thymine (T) (T-to-C conversion) upon reverse transcription (Herzog et al. 2017). The chemical conversion greatly reduces the labor associated with multi-time point metabolic labeling experiments by eliminating the need for biochemical enrichment techniques and allowing the use of a lower starting amount for construction of cDNA libraries. Moreover, modeling of RNA decay is simplified due to the elimination of accounting for biotin conjugation and separation of biotinylated RNAs efficiency in the modeling parameters.

SLAM-Seq has been utilized in mammalian and *Drosophila* cells lines using 4sU (Herzog et al. 2017; Muhar et al. 2018; Reichholf et al. 2019), however, no reports exist for the use for SLAM-Seq in yeast. We reasoned that SLAM-Seq would have a distinct advantage over transcriptional inhibition methods due to its minimally invasive nature, and would be less laborious that currently used pull-down methods. Given that this method can be easily utilized to measure RNA half-life, we measured the RNA decay in the BY4741 wt strain and the isogenic *upf3Δ* mutant. The aim was to identify NMD substrates since half-life measurements are the golden standard for this as opposed to just measuring changes in steady-state levels in the transcriptome (Fig. 1).

**Figure 1.**
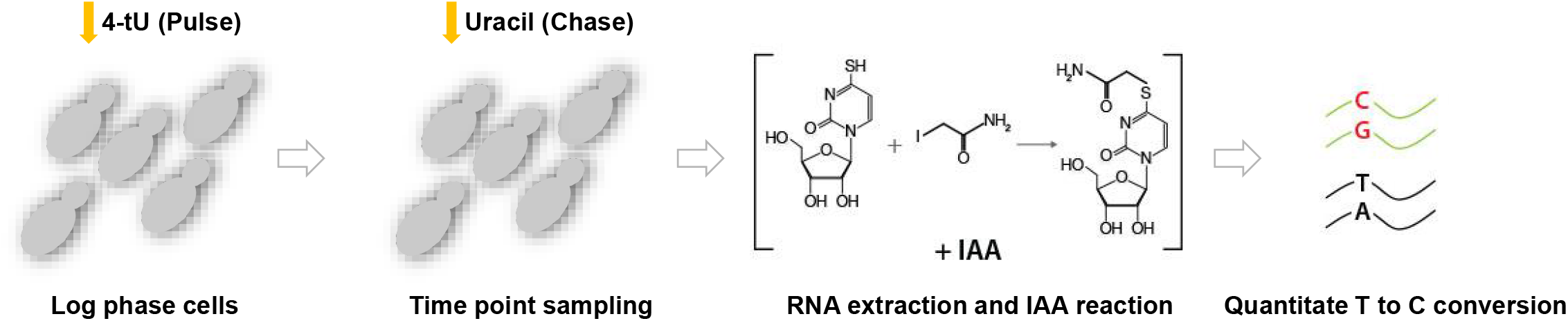
SLAM-Seq work flow Log phase cells are labeled with 4-tU followed by chasing the label with a high concentration of unmodified uracil. 4-tU is taken up by cells and incorporated into RNA. Samples are taken during the chase and metabolically inactivated using chilled ethanol. RNA is then extracted and incorporated 4-tU is alkylated with iodoacetamide (IAA), yielding a stable derivative of the thiol-containing 4-tU. Reverse transcriptase misincorporates a guanine rather than adenine at derivatized positions during library construction, causing a T to C conversion. These are then quantitated using the SLAMDUNK pipeline.

### Confidence of half-life estimates and coverage of RNA species using SLAM-Seq in yeast

We started by treating cells in logarithmic growth phase with a low concentration of 4tU for 2 h followed by chasing the labeled transcripts with a high concentration of uracil. The samples were incubated for 2 min prior to the onset of sampling to account for the inefficient chase that has been previously observed (Chan et al. 2018) and time points were sampled at 0, 1, 4, 8, 18, and 58 min. Subsequently RNA was extracted, alkylated, reverse transcribed, and sequenced. We observed a T-to-C conversion rate ranging between 2.54 % and 2.83 % for BY4741 at time point 0 and between 2.41 % and 2.55 % in *upf3Δ*, similar to observations in mouse embryonic stem cells (Herzog et al. 2017) with the conversion rate decreasing with increasing chase time as expected (see Materials and Methods for details and Supplementary files S1-S4 for quality control reports and sample indices). We started by comparing the performance of SLAM-Seq to other methods. We limited the analysis to transcripts with a measureable half-life in all three biological replicates with a coefficient of variation less than 0.35 (Herzog et al. 2017). Full unfiltered dataset can be found in Supplementary file S5.

Using all six time points, we were able to fit 4049 transcripts in BY4741 that meet the criteria above representing 67.5 % of all annotated ORFs in the wt or 72.8 % of the ORFs available in our 3’-UTR annotation, and 3638 transcripts in *upf3Δ* (60.7 % of all annotated ORFs and 65.4 % of ORFs available in our 3’-UTR annotation). The linear distribution of the half-life was right-skewed as expected (Fig. 2 A). The wt BY4741 strain showed a sharper peak than *upf3Δ*, with the individual biological replicates showing high correlation coefficients, ranging between 0.88 and 0.93 in BY4741 and between 0.83 and 0.88 in *upf3Δ* (Fig. 2 B). Gene Ontology (GO) term enrichment analysis of the 1 % most long-lived transcripts indicated that those encoding proteins involved in housekeeping functions such as glycolysis and translation had the longest half-life (Fig. 2 A; a full enrichment table is in Supplemental file S6), while the 1 % with the shortest half-life did not show any particular enrichment.

**Figure 2.**
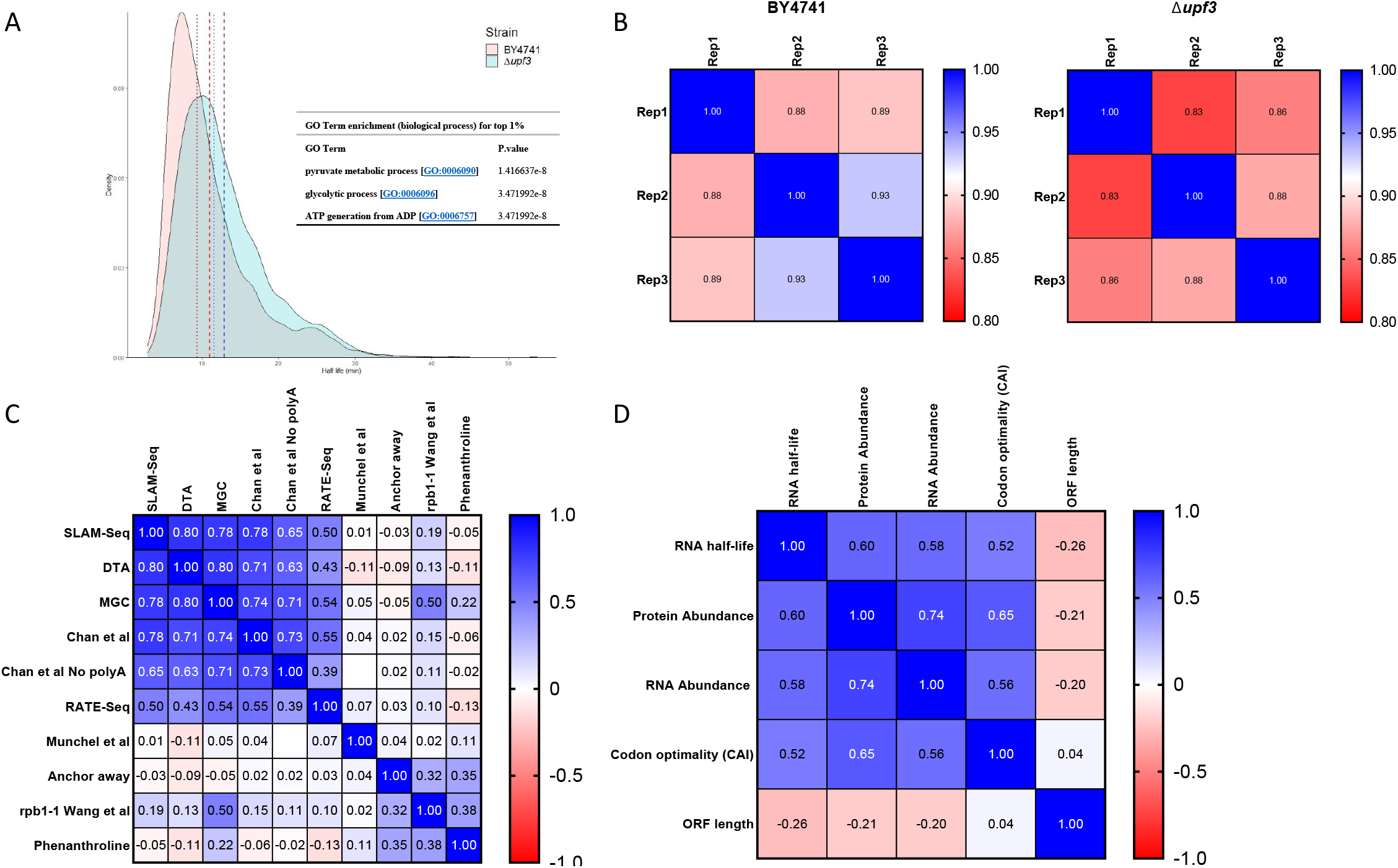
Half-life distribution and comparison to other methods **A)** Half-life distribution in wt and *upf3Δ* The dashed line indicates mean and dotted line the median; red is for wt and blue for *upf3Δ*. Partial enrichment analysis for the longest lived 1 % of the transcripts is shown as an inset (full GO term enrichment for longest lived 1 % is listed in Supplementary Table S6). **B)** Inter-replicate correlation using Pearson’s correlation coefficient **C)** Comparison of half-lives calculated using SLAM-Seq to other methods using Spearman’s Rho DTA (Miller et al. 2011), MGC (Baudrimont et al. 2017; Chan et al. 2018), RATE-Seq (Munchel et al. 2011; Neymotin et al. 2014), Anchor away (Geisberg et al. 2014), *rpb1-1* (Wang et al. 2002), phenanthroline (Grigull et al. 2004). **D)** Correlation of half-lives to other transcripts features using Spearman’s Rho RNA abundance was calculated using the mean CPM across all the time points of wt, protein abundance was from Ho et al. (2018), codon optimality was from Drummond et al. (2006), and ORF length from annotations of *S. cerevisiae* S288C (assembly R64).

### Correlation of SLAM-Seq with other methods for mRNA turnover measurements

Next, we compared SLAM-Seq and other mRNA decay measurements using halflives calculated from the BY4741 strain. Fig. 2 C shows the Spearman correlation between various studies employing transcriptional inhibition, transcriptional shutoff or metabolic labeling. As expected, transcriptional inhibition methods correlated very poorly with each other and with other methods, while multiplexed gene control and variants of metabolic labeling correlated well with each other and with SLAM-Seq, the rho values ranging between 0.5 and 0.8. It has been previously suggested that a Spearman rho value of 0.7 and above indicates high inter-method reliability (Wada and Becskei 2017). We observe high correlation (0.78) with multiplexed gene control (MGC), the metabolic labeling study with improved RNA separation (Chan et al. 2018); 0.78, and the Dynamic Transcriptome Analysis (DTA) method (Miller et al. 2011) (0.8), indicating that SLAM-Seq is able to classify transcripts as stable or unstable in good agreement with other recent methods. However, the absolute magnitude of measured half-life is different between the methods. Chan et al. (2018) and studies using MGC calculate shorter half-life than DTA and SLAM-Seq (see Discussion). Finally, we checked for half-life correlation with other ORF features (Fig. 2 D). The results mirrored that of Chan et al. (2018), namely a weak inverse correlation of mRNA half-life with ORF length (rho = −0.26) and a positive correlation with protein abundance (rho = 0.6), RNA abundance (rho = 0.58) and codon optimality (rho = 0.52) all which indicates that highly expressed transcripts and transcripts with optimal codons tend to have longer halflives.

### Assignment of transcripts as NMD targets

We found 3033 transcripts passing the above filtration parameters that were in common between *upf3Δ*, and could be directly compared. We found 346 transcripts to have a significantly altered half-life in *upf3Δ* (q value of < 0.05 and half-life fold change of at least 1.5). However, the criteria used to filter the transcripts for comparison was too restrictive for reporting NMD targets, since if two replicates reported similar half-life in wt but the other replicate did not pass the fitting filter (R^2^ ≥ 0.6) and the replicates in *upf3Δ* passed all the other criteria, the transcript would still be excluded from the analysis due to the lack of one half-life measurement. Therefore, we used ≥ 2-fold half-life fold change as criteria for the other transcripts that were not part of the previously compared 3033 transcripts and had at least two measurements in one strain and three measurements in the other. Combining the above analyses, we found a total of 598 transcripts with altered half-life between the two strains (full list can be found in Supplementary file S7). Additional transcripts that had less than 1 count per million (CPM) in any of the time points and hence filtered out in the wt even though a number of them had increased read counts in *upf3Δ*, were not considered in this set since we could not calculate their half-lives. These transcripts, thus representing weakly expressed mRNAs only detectable in *upf3Δ* mutants. The transcripts that failed the CPM filtration in wt can be found in Supplementary Table S8. The majority of the transcripts with an altered half-life relative to wt had an increase half-life in *upf3Δ* (n = 580) with only 18 transcripts showing a decreased half-life in *upf3Δ* and have no particular enrichment. Considering that the half-life of NMD substrates increases rather than decreases, we only considered the transcripts with increased half-life ratio for further analysis.

We were able to recover transcripts that represent the common structural classes regulated by NMD (Fig. 3): 1) non-sense codon containing *YML002W*. In our strain background, *YML002W* is continuous with *YML003W* and a single thymidine deletion in *YML003W* causes a frameshift and a premature stop codon hence rendering the transcript sensitive to NMD; 2) mRNAs utilizing frameshifting in their translation (*EST3*); 3) uORF-containing transcripts (*CPA1*); 4) transcripts with long 3’-UTR (*COX19*). Moreover, we recover 8 additional transcripts that were previously identified in low-throughput studies (Fig. 4). We compared our identified NMD targets with previous studies (Table 1) and found an overall overlap of 60 %, indicating that we were able to successfully identify NMD substrates based on changes in half-life. The group of targets were analyzed for enrichment of GO terms, and were found to be enriched for two subgroups of interest: protein sumoylation by Molecular function, and spliceosome by Cellular component (Table 2).

**Figure 3.**
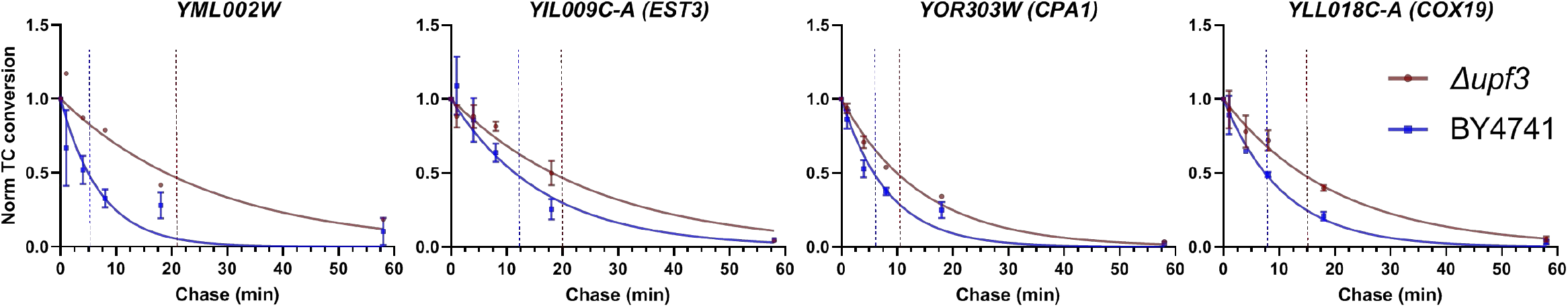
Transcripts representing common structural classes known to be regulated by NMD Background subtracted and chase onset normalized T-to-C conversion rates were fitted to first order exponential decay. *YML002W:* non-sense codon containing in this strain background *EST3:* translational frame-shifting *CPA1:* uORF-containing *COX19:* long 3’-UTR Error bars represent the standard deviation. Error bars for *YML002* in *upf3Δ* were not added since it was calculated from two replicates.

**Figure 4.**
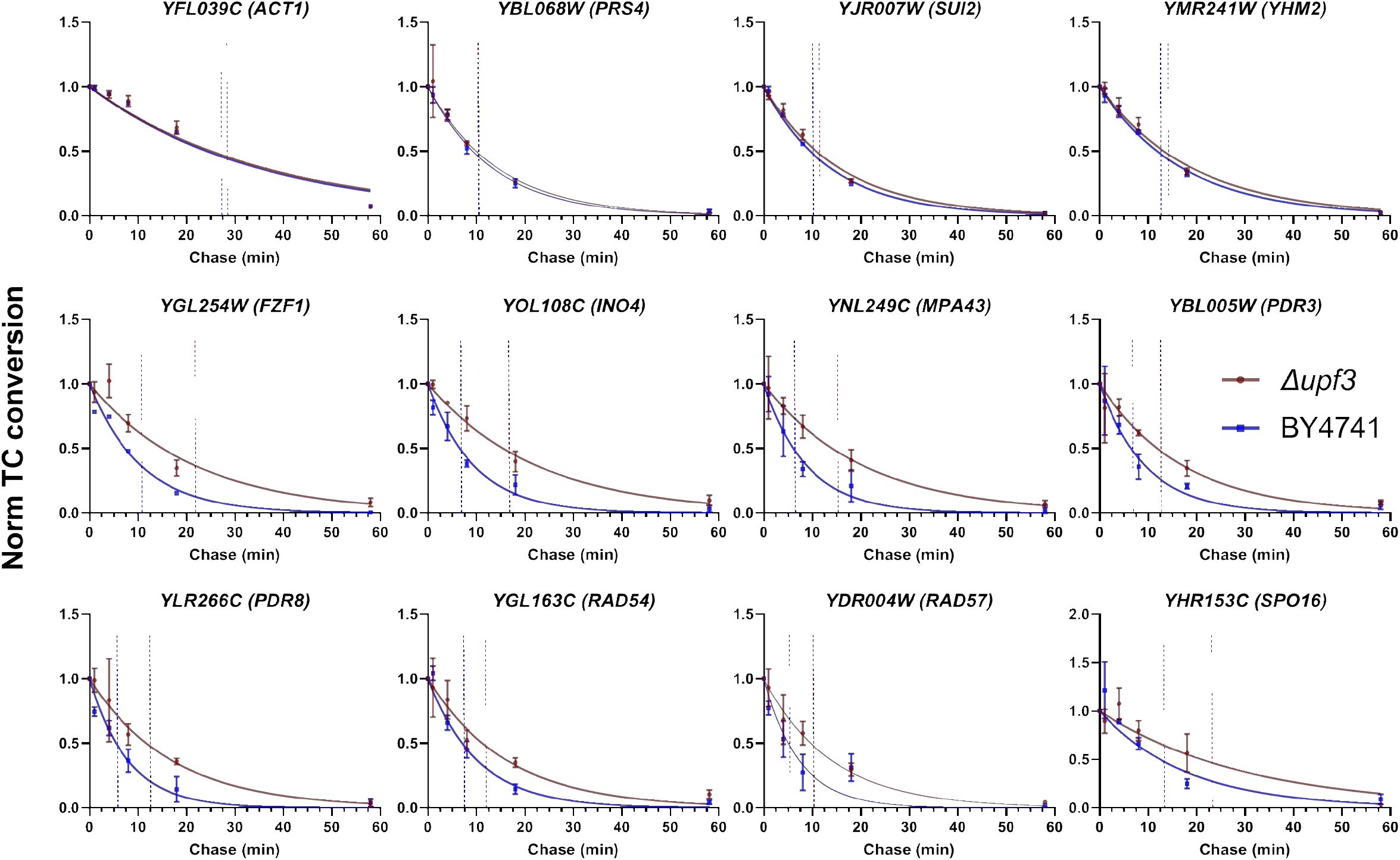
Transcripts found to be NMD regulated in previous low throughput studies Background-subtracted and chase onset normalized T-to-C conversion rates were fitted to first order exponential decay. The first row is control transcripts not regulated by NMD. Error bars represent standard deviation. Transcripts with no error bars were calculated from two replicates. Transcripts were found to be NMD regulated in the following studies: *FZF1, INO4, PDR3* and *PDR8* (Guan et al. 2006); *RAD54* and *RAD57* (Janke et al. 2016); *MPA43* (Kebaara and Atkin 2009); *SPO16* (Zaborske et al. 2013).

**Table 1.** Overlap with other studies aiming to identify NMD targets

| Overlap with other studies |  |  |  |  |  |
| --- | --- | --- | --- | --- | --- |
|  | He et al. (2003) | Johansson et al. (2007) | Johansson et al. (2007) | Malabat et al. (2015) | Celik et al. (2017) |
| Method | Microarray | UPF1-TAP affinity | NMD reactivation | RNA-Seq | RNA-Seq |
| Common hits | 189 | 147 | 157 | 242 | 291 |
| Percentage of SLAM-Seq positives found | 32.6 | 25.3 | 27.0 | 41.6 | 50.0 |

Number of common hits and percentage of SLAM-Seq hits in this study found by other studies. He et al. (2003) used microarrays to detect upregulated transcripts in NMD defective strains. Johansson et al. (2007) tested for enrichment of NMD target binding to Upf1 using tandem affinity purification and for transcripts downregulated when *GALI*-driven expression of *UPF2* is activated. Celik et al. (2017) and Malabat et al. (2015) used RNA-Seq to determine differential abundance between wt and NMD defective cells.

**Table 2.** Enrichment analysis for transcripts with increased half-life in *upf3Δ*

| **GO Term enrichment (biological function) for NMD hits** |  |
| --- | --- |
| **GO Term** | **P value** |
| regulation of protein sumoylation [GO:0033233] | 0.026628 |
| positive regulation of protein sumoylation [GO:0033235] | 0.026628 |
| **GO Term enrichment (cellular component) for NMD hits** |  |
| **GO Term** | **P value** |
| spliceosomal complex [GO:0005681] | 0.002926 |
| nuclear protein-containing complex [GO:0140513] | 0.005249 |
| intracellular membrane-bounded organelle [GO:0043231] | 0.019221 |
| membrane-bounded organelle [GO:0043227] | 0.022166 |
| membrane [GO:0016020] | 0.037183 |

## DISCUSSION

Recent understanding and chemical derivatization of RNA species has led to the emergence of new techniques that simplify the measurements of previously difficult to capture parameters such as RNA-half life on a genome wide scale. SLAM-Seq has been heavily used in mammalian cells (Baptista and Dölken 2018; Matsushima et al. 2018; Muhar et al. 2018), however, no reports for simpler eukaryotic organisms are available. In this work, we conducted an assessment of the applicability to SLAM-Seq to *S. cerevisiae*, and found excellent correlation with recent studies utilizing metabolic labeling. Moreover, we applied SLAM-Seq to measure changes in RNA half-lives in NMD defective cells, and were able to identify NMD substrates based on actual RNA half-life change as opposed to increased steady-state RNA levels. Using SLAM-Seq, we could avoid the problems associated with other techniques such as transcriptional inhibition (*e.g. rpb1-1*), which essentially measures RNA half-life in dying cells, hence giving rise to physiologically irrelevant half-life measurements that are poorly reproducible, and also bypassed the requirement for ample starting material and spike-in normalization required for pull-down approaches. Using SLAM-Seq we were able to calculate half-lives with high intersample and intermethod correlation as opposed to the poorly correlating values reported previously in transcriptional inhibitions studies due to the problems highlighted above.

Slam-Seq was paired using 3’-end mRNA sequencing (QuantSeq), which alleviates the cost of full RNA seq, especially given the multi-timepoint experiments that are required for half-life determination, and also facilitates downstream analysis due to the lack of a requirement for transcript length normalization. The QuantSeq protocol captures fully processed RNA pol II transcripts, which can be advantageous if that is the subset of interest. However, the inability to distinguish RNA isoforms may pose a problem for certain applications. In our particular case, we were unable to detect transcripts belonging to an NMD class consisting of inefficiently spliced transcripts that leak to the cytoplasm (Celik et al. 2017) with the exception of *HRB1*. This could indicate that the either the mature form of *HRB1* could also be under NMD regulation, or that since *HRB1* has lower expression compared to other transcripts belonging to the same class (*e.g. YBR089C-A, YFL034C-A* and *YGR148C*) the increase in intron-containing unspliced isoforms can be detectable with halflife change due it being able to compete with the mature form for labeling and hence can contribute to half-life calculation while for the other more strongly expressed transcripts, this effect is too low to be seen. Nonetheless, this does not detract from the value of the technique as *S. cerevisiae* is intron poor and this problem can be remedied with other types of RNA-Seq libraries as long as they incorporate a reverse transcription step (Herzog et al. 2017).

We found a high correlation in the classification of transcripts into stable and unstable transcript with other metabolic labeling studies. However, the absolute value for RNA halflife was not consistent between the studies. This has been an ongoing problem for half-life determination in yeast due to earlier reports indicating a higher half-life value but considering that earlier studies utilized transcriptional inhibition, it is not surprising to see higher half-life values that poorly correlate with our study and even other studies using derivatives of transcriptional inhibition. In this work we report mean half-life value of 11 min (median 9.4) which agrees more closely with Miller et al. (2011) in which a mean value of 14 min (median 11 min) was reported, while the other high correlating studies (Baudrimont et al. 2017; Chan et al. 2018) reported much lower value than all previous measurements. These differences might be attributable to methodic difference; an earlier study (Munchel et al. 2011) which had lower separation efficiency of labeled transcripts compared to the second generation assay (Chan et al. 2018), calculated values that did not correlate well with any of the previous studies. This highlights the technical difficulties associated with pull-down approaches and the effect of the pull-down efficiency on half-life calculation. Baudrimont et al. (2017) similarly reported a lower half-life mean than previous estimates agreeing more closely to that of Chan et al. (2018). It has been argued that slower than expected kinetics of nucleotide incorporation might be the reason for these discrepancies, since in MGC the target transcript in placed under a modified controllable promoter and decay is measured post-shutoff, and the delay from nucleotide incorporation is eliminated (Wada and Becskei 2017). However, Chan et al. (2018) used a pulse-chase approach and still calculated values similar to those of Baudrimont et al. (2017). Regardless of the difference in the absolute values the high correlation of the methods (hence ability to appropriately group transcripts as stable and unstable) can be appreciated since all those techniques are methodically independent. Moreover, if indeed the delayed incorporation kinetics has an effect on half-life calculation future correction can be made using a factor calculated based on Baudrimont et al. (2017) given that data for more transcripts becomes available.

We report new NMD substrates based on changes in RNA half-life, rather than steady-state abundance, and were able to identify 230 new transcripts, whereas the remaining 60 % overlapped with previous studies (Table 1). We find it likely that a large fraction of the 230 newly identified NMD targets in this work, using altered decay rates in *upf3Δ* mutants as criterion, escaped detection in earlier studies using mRNA steady-state levels. The highest overlap with a single previous study (50 %) was with Celik et al. (2017), where RNA-Seq was utilized to identify NMD targets in high resolution, indicating that SLAM-Seq agrees well with more recent datasets that aimed to identify NMD targets with the added benefit of producing a quantitative value for the half-lives. We successfully capture major classes of transcripts expected to be NMD targets, with the exception of the inefficiently spliced transcripts class (see above). We found an enrichment for spliceosomal factors in agreement with what has been found in mammalian cells (Saltzman et al. 2008). We envision that SLAM-Seq would be highly useful for studies looking to identify changes in transcripts stabilities under various condition such as those looking at the effect of particular drugs on the cells or metabolic changes under different nutritional conditions such as nitrogen sources (Miller et al. 2018) in addition to the application to NMD pathway substrate detection as we describe here.

## MATERIALS AND METHODS

### Strains and media

BY4741 (MATa *his3Δ1 leu2Δ0 met15Δ0 ura3Δ0*) was used as the control strain (WT) and an isogenic *upf3Δ* strain was used as the tester. YPD (2 % glucose, 2 % peptone, and 1 % yeast extract) was used for routine culturing of strains. Low uracil medium (LUM; 0.19 % yeast nitrogen base, 0.5 % ammonium sulfate, 2 % glucose and 0.077 % Complete Supplement Mixture without uracil (-URA CSM; ForMedium), supplemented with 0.1 mM uracil) was used for labeling conditions. The chase medium used to remove the RNA label was identical to LUM except being supplemented with 20 mM uracil. Liquid cultures were grown in a rotary shaker at 30°C at 200□rpm.

### 4tU metabolic labeling

Overnight cultures were made in LUM and diluted to OD_600 nm_ = 0.1 on the next day in a total volume of 200 ml, then grown to OD_600 nm_ = 0.4. Subsequently the cultures were split into two 100 ml cultures and 4tU (Sigma) was added to one culture to a final concentration of 0.2 mM and DMSO was added to the negative control culture to final concentration of 0.02 %. Both cultures were then grown for an additional 2 h. To chase the 4tU label, the cultures were then briefly centrifuged, the 4tU-containing media removed and replaced with an equivalent volume of chase medium and incubated for 2 min at the same growth condition used for labeling. Time 0 was taken after the 2 min incubation with the chase media and at 1, 4, 8, 18, and 58 min after time 0. For each time point, a 10 ml aliquot was taken and added to an equal volume of 95 % ethanol pre-chilled at −70°C, and then kept on dry ice until the end of the sampling. Only one time point was sampled for the DMSO-treated cultures to calculate the background T-to-C conversion rate. This treatment was repeated in biological triplicates for both WT and *upf3Δ*.

### RNA extraction

Total RNA was extracted from the sampled cell pellets using Quick-RNA Fungal/Bacterial Kit (Zymo Research) as per manufacturer instructions with the modification of adding Reducing agent (RA) from the SLAM-Seq Kinetics Kit - Catabolic Kinetics Module (Lexogen) to all extraction reagents of the RNA extraction kit as indicated by the manufacturer. Subsequently the RNA was alkylated and purified according to the instruction of the SLAM-Seq Kinetics Kit - Catabolic Kinetics Module.

### Library preparation

Libraries were prepared using the QuantSeq 3’ mRNA-Seq Library Prep Kit FWD for Illumina (Lexogen). Library preparation, sequencing in SR75 mode, and read demultiplexing was carried out by Lexogen Services.

### Bioinformatics analysis and half-life analysis

Demultiplexed reads were analyzed using SLAM DUNK (https://t-neumann.github.io/slamdunk/) (Neumann et al. 2019) through the nf core SLAM-Seq processing and analysis pipeline (Ewels et al. 2020). Default settings were used with the exception of designating the read length manually. A custom BED file containing the position of the ORFs 3’-UTRs was created based on three annotations (Xu et al. 2009; Roy and Chanfreau 2020) and our own analysis of the TIF-seq data described previously (Xu et al. 2009; Pelechano et al. 2013) and used for defining the windows for 3’-UTRs (Supplementary files S9 and S10). The resulting count files were filtered to include genes that had at least 1 CPM in their corresponding time series. Subsequently T-to-C conversion rates were corrected by subtraction of the background conversion rate of the corresponding DMSO-treated samples and normalized to the onset of the chase (time 0). Half-lives were calculated using the minpack.lm package as described previously (Herzog et al. 2017) and values that had an R^2^ value of less than 0.6 were filtered out from the resulting calculated half-lives, exact sampling times were used to reduce error associated with slight deviations between theoretical and actual sampling time points, with actual time points being the moment the sample is mixed with the chilled ethanol (see Supplementary file S11 for exact time points). Values of half-lives were corrected for growth during the chase using the halflife formula in Munchel et al. (2011). Transcripts that passed the more stringent criteria n = 3033 (all replicates had values and coefficient of variation less than 0.35) were tested using unpaired t test with single pooled variance assumption after log_10_ transformation of the values. Transcripts with a q value of less than 0.05 and a fold change in half-life of 1.5 or more where considered positives. For transcripts that were not part of the above set, a ≥ 2-fold half-life fold change was used as a criteria and each transcript needed to have at least two measurements in one strain and three measurements in the other. Enrichment analysis was carried out using Yeastmine (https://yeastmine.yeastgenome.org/yeastmine/). Graphing of RNA degradation curves was done using GraphPad Prism 9.

## ACKNOWLEDGEMENTS

We thank Vicent Pelechano for providing Xu’s transcriptome annotation (Xu et al. 2009) and Allan Jacobson for providing datasets from He et al. (2003). This work was supported by a grant from the Swedish Cancer Fund (19 0133).

## SUPPLEMENTARY FILES

**Supplementary File S1** Quality check report, experimental replicate 1

**Supplementary File S2** Quality check report, experimental replicate 2

**Supplementary File S3** Quality check report, experimental replicate 3

**Supplementary File S4** Index for all samples, all replicates

**Supplementary File S5** Full unfiltered dataset for both strains

**Supplementary File S6** GO term enrichment for the 1 % most long-lived transcripts

**Supplementary File S7** Full list of transcripts with altered half-life in *upf3Δ* mutants

**Supplementary File S8** Transcripts with too few counts in wt to be considered for quantitative evaluation, even if their number of counts was increased in *upf3Δ* mutants

**Supplementary File S9** Description of construction of 3’-UTR reading windows

**Supplementary File S10** BED file of all 3’-UTR reading windows

**Supplementary File S11** Exact sampling times

